## Supplementary material for "The microbiota of *Amblyomma americanum* reflects known westward expansion": R script

# Tick Microbiome analysis
# Target region V4-V5
getwd()
setwd("~/Desktop/TickData/")

packageVersion("dada2")  #1.16.0

#Specify path where FASTQ files are located
pathT <- "~/Desktop/TickData/"
pathT
list.files(pathT)

#Sort files to ensure forward/reverse reads are in the same order
# Forward and reverse fastq filenames have format: SAMPLENAME_R1_001.fastq and SAMPLENAME_R2_001.fastq
fnFsT <- sort(list.files(pathT, pattern = "_R1_001.fastq.gz",full.names = TRUE))
fnFsT
fnRsT <- sort(list.files(pathT, pattern = "_R2_001.fastq.gz",full.names = TRUE))
fnRsT

# Extract sample names, assuming filenames have format: SAMPLENAME_XXX.fastq
#sample.namesT <- sapply(strsplit(fnFsT, "_"), `[`,1)
#sample.namesT

## Ran into a problem when filtering an triming downstream as samples are not named
# with a homogeneous system. Some samples have leters to discriminate between the same collection site
# others have "_" followed by a number. This interferes with the name extraction later on as some samples end up with the same name
# as they drop the discriminant number after the undescore...

sample.namesT2 <- sapply(strsplit(basename(fnFsT), "_"), `[`, 1)
sample.namesT2  #190 samples

# Therefore we can recur to the second field after the "_" to take back the discriminant
# names become larger but we are able to individualize each of the 190 samples
sample.fields <- lapply(strsplit(basename(fnFsT), "_"), `[`, c(1,2))
sample.namesT <- sapply(sample.fields, function(fields) paste0(fields[1], fields[2]))
sample.namesT


#Inspect read quality profiles
plotQualityProfile(fnFsT[1:2])
plotQualityProfile(fnFsT[24:25])
plotQualityProfile(fnFsT[189:190]) 
# do this for samples and the blank
plotQualityProfile(fnRsT[1:2])
plotQualityProfile(fnRsT[24:25])

### As the project was executed with a seq strategy that will not allow us to succesfully overlap Fs and Rs reads
# we will proceed only with Forward reads as sugested by the developer and several other sources. There is an alternative
# to concatenate them at the merge pairs stage but its not encouraged.
## DADA2 forum link referring to this issue and why to proceed with F's for this hypervariable region (V4):https://forum.qiime2.org/t/using-dada2-losing-50-of-reads-after-filtering-and-80-of-reads-by-the-end/12745/4


# Place filtered files in filtered/ subdirectory

filtFsT <- file.path(pathT, "filtered", paste0(sample.namesT, "_F_filt.fastq.gz"))
filtRsT <- file.path(pathT, "filtered", paste0(sample.namesT, "_R_filt.fastq.gz"))
names(filtFsT) <- sample.namesT
names(filtRsT) <- sample.namesT
filtFsT
filtRsT

#Verifying if the solution of taking the sencond element after "_" in the full file names
# resolved the problem of duplicated names downstream
any(duplicated(sample.namesT))
any(duplicated(c(fnFsT, fnRsT)))
any(duplicated(c(filtFsT, filtRsT)))   ## ALL FALSE so no duplicated names


outT <- filterAndTrim(fnFsT, filtFsT, fnRsT, filtRsT, truncLen=c(150,150),
                      maxN=0, maxEE=c(2,2), truncQ=2, rm.phix=TRUE,
                      compress=TRUE, multithread=TRUE) 
head(outT)

#reads.in reads.out
#AL129127A_S143_L001_R1_001.fastq.gz     6324      6187
#AL129127B_S144_L001_R1_001.fastq.gz    45998     45379
#AL129127C_S145_L001_R1_001.fastq.gz    24990     24757
#AL129127D_S146_L001_R1_001.fastq.gz    34754     34361
#AL129127E_S147_L001_R1_001.fastq.gz    33942     33537
#AL129127F_S148_L001_R1_001.fastq.gz    28151     27789

# Learning the error rates
errFT <- learnErrors(filtFsT, multithread=TRUE)
# errRT <- learnErrors(filtRsT, multithread=TRUE)  done ONLY for F's

## Ran in 1 minute connected to the server for each pair
# 103655250 total bases in 691035 reads from 21 samples will be used for 
# learning the error rates.

#visualize the estimated error rates
plotErrors(errFT, nominalQ=TRUE)
#  ---> every plot 
# Here the estimated error rates (black line) are a good fit to the observed rates 
# (points), and the error rates drop with increased quality as expected. Everything looks 
# reasonable and we proceed with confidence.

# IMPORTANT! Proceeding only with forward reads as overlap is doubtful and concatenating is not 
# encouraged for ASV analyses--> Even if dada2 can handle the spacers it would affect the Add species step among other issues
dadaFsT <- dada(filtFsT, err=errFT, multithread=TRUE)
dadaFsT[[1]]

# source / forum suggesting using V4 F's only:https://forum.qiime2.org/t/using-dada2-losing-50-of-reads-after-filtering-and-80-of-reads-by-the-end/12745/4

# dada-class: object describing DADA2 denoising results
# 18 sequence variants were inferred from 836 input unique sequences.
# Key parameters: OMEGA_A = 1e-40, OMEGA_C = 1e-40, BAND_SIZE = 16


# Pooling information across samples can increase sensitivity to sequence variants that may be present
# at very low frequencies in multiple samples. The dada2 package offers two types of pooling. dada(..., pool=TRUE) 
# performs standard pooled processing, in which all samples are pooled together for sample inference. 

dadaFsT2 <- dada(filtFsT, err=errFT, multithread=TRUE, pool=TRUE)
# 190 samples pooled: 6753479 reads in 530350 unique sequences.
dadaFsT2[[1]]
# dada-class: object describing DADA2 denoising results
# 106 sequence variants were inferred from 836 input unique sequences.
# Key parameters: OMEGA_A = 1e-40, OMEGA_C = 1e-40, BAND_SIZE = 16


# NO MERGING since we are working only with forward reads
# Construct sequence tab
seqtabT <- makeSequenceTable(dadaFsT2)
dim(seqtabT)  #[1]  190 5632

# Inspect distribution of sequence lengths
table(nchar(getSequences(seqtabT)))
# 150bp --> 5632  # all ASVs have the expected lenght, 
# but smaller than the lenght to cover all of of the targeted V4 region...

# Chimera removal
seqtab.nochimT <- removeBimeraDenovo(seqtabT, method="consensus", multithread=TRUE, verbose=TRUE)
# Identified 221 bimeras out of 5632 input sequences.
dim(seqtab.nochimT)
# [1]  190 5411   --> 96% sequence variants remained
sum(seqtab.nochimT)/sum(seqtabT)  # --> [1] 0.9935885 --> chimeric variants account for less than 1% of the reads  


getN <- function(x) sum(getUniques(x))
track <- cbind(outT, sapply(dadaFsT2, getN),rowSums(seqtab.nochimT))
colnames(track) <- c("input", "filtered", "denoisedF","nonchim")
rownames(track) <- sample.namesT
head(track)
#we can verify the samples at the end of the list
tail(track)

#How does our blank sample looks like?
rownames(track)[25]
track[25,] # Blank sample data

#Head
#              input filtered denoisedF nonchim
#AL129127AS143  6324     6187      6117    6098
#AL129127BS144 45998    45379     45355   45309
#AL129127CS145 24990    24757     24742   24737
#AL129127DS146 34754    34361     34349   34296
#AL129127ES147 33942    33537     33516   33511
#AL129127FS148 28151    27789     27783   27764

#     Blank1  input  filtered denoisedF   nonchim 
#              2349      2282      2251      2249 


# Looks good! We kept the majority of our raw Forward reads, 
# and there is no over-large drop associated with any single step.

### TAXONOMY assignation using the latest Silva training set

taxaT <- assignTaxonomy(seqtab.nochimT, "~/Desktop/TickData/silva_nr_v132_train_set.fa.gz",multithread=TRUE)
taxaT <- addSpecies(taxaT, "~/Desktop/TickData/silva_species_assignment_v132.fa.gz")

taxa.printT <- taxaT # Removing sequence rownames for display only
rownames(taxa.printT) <- NULL
head(taxa.printT)
tail(taxa.printT) ## Many Eukaryota NA NA NA... classifications. 
# reads may be in the opposite orientation as the reference database

## Doing Tax assignation with tryRC=TRUE to search the DB using the reverse complement
taxaT <- assignTaxonomy(seqtab.nochimT, "~/Desktop/TickData/silva_nr_v132_train_set.fa.gz",tryRC=TRUE,multithread=TRUE)
taxaT <- addSpecies(taxaT, "~/Desktop/TickData/silva_species_assignment_v132.fa.gz")

taxa.printT <- taxaT # Removing sequence rownames for display only
rownames(taxa.printT) <- NULL
head(taxa.printT)
tail(taxa.printT)
## Still we have Eukaryota NA NA NA... classifications in the tax table.
## we will have to filter them out from the Phyloseq objetc downstream along with mitochondria and chloroplast elements


## Note: We could use DECIPHER to make the tax assign step too... its performance and speed are better. 

#######   PHYLOSEQ   ###########

SampledataTicks<-read.csv("~/Desktop/TickData/Mapping_file_Microbiome_Nov19_Aamericanum.csv", header = TRUE, sep = ";", row.names=1)
SampledataTicks
class(SampledataTicks)

#ordering the sample data data.frame by ID alphabetical order 
SampledataTicks<-SampledataTicks[order(rownames(SampledataTicks)), ]
sampleIDoriginal<-rownames(SampledataTicks)
sampleIDoriginal ## saving the names without the character string attached upstream as a solution for non homogeneity in the coding system
sample.namesT

colnames(SampledataTicks)
SampledataTicks<-SampledataTicks[,c(5,6,14,15,16,17)]
SampledataTicks
rownames(SampledataTicks)<-sample.namesT
SampledataTicks$Type<-factor(c(rep("Sample",24),rep("Blank",1),rep("Sample",165)))
SampledataTicks$Sex<-factor(SampledataTicks$Sex)
SampledataTicks

# creating Phyloseq object: OTU table, Tax table, Sample ID, Phylogeny, ref seq

library(phyloseq); packageVersion("phyloseq")
library(Biostrings); packageVersion("Biostrings")
library(ggplot2); packageVersion("ggplot2")

#Extracting the standard goods from R
# giving the seq headers more manageable names (ASV_1, ASV_2...)
asv_seqsT <- colnames(seqtab.nochimT)
asv_seqsT

# The dimensions of our matrix are 52 x 429
asv_headersT <- vector(dim(seqtab.nochimT)[2], mode="character")  # vector with spaces corresponding to ASV names
asv_headersT

for (i in 1:dim(seqtab.nochimT)[2]) {
  asv_headersT[i] <- paste(">ASV", i, sep="_")
}

asv_headersT  # Vector with labels for the true ASVs detected, whose sequences are on asv_seqsT variable

# making and writing out a fasta of the final ASV seqs:
asv_fastaT <- c(rbind(asv_headersT, asv_seqsT))
asv_fastaT
write(asv_fastaT, "ASVs_Tickmicrobiota.fa")  #fasta file 


# count table:
asv_tabT <- t(seqtab.nochimT)
asv_tabT
row.names(asv_tabT) <- sub(">", "", asv_headersT)
write.table(asv_tabT, "ASVs_countsTickmicrobiota.txt", sep="\t", quote=T)
asv_tabT

ASVcountsT <-t(asv_tabT)
ASVcountsT  # transposed and with ASV_ structure to build the phyloseq object compatible with our explanatory matrix: 
# samples as rows.

# tax table:
asv_taxT <- taxaT
row.names(asv_taxT) <- sub(">", "", asv_headersT)
asv_taxT
write.table(asv_taxT, "ASVs_taxonomyTickmicrobiota.txt", sep="\t", quote=T)
## including Eukaryotes and NA's.... Eukaryotic parasites perhaps? Remember we used only 150bp Fw reads to assign tax...resolution will be hampered


## Assembling the Phyloseq object

TickPS <- phyloseq(otu_table(seqtab.nochimT, taxa_are_rows=FALSE), 
                   sample_data(SampledataTicks), 
                   tax_table(taxaT))

TickPS

# Merging phyloseq object with ASV sequences
dnaT <- Biostrings::DNAStringSet(taxa_names(TickPS))
names(dnaT) <- taxa_names(TickPS)
TickPS <- merge_phyloseq(TickPS, dnaT)
taxa_names(TickPS) <- paste0("ASV", seq(ntaxa(TickPS)))
TickPS
# DNA sequences of our ASVs are now stored in the refseq slot of the phyloseq object, 
# and taxa were renamed with a shorter string --> "ASVX

# phyloseq-class experiment-level object
# otu_table()   OTU Table:         [ 5411 taxa and 190 samples ]
# sample_data() Sample Data:       [ 190 samples by 7 sample variables ]
# tax_table()   Taxonomy Table:    [ 5411 taxa by 7 taxonomic ranks ]
# refseq()      DNAStringSet:      [ 5411 reference sequences ]
tax_table(TickPS)[1,]

# Example ASV1 

#    Taxonomy Table:     [1 taxa by 7 taxonomic ranks]:
#        Kingdom    Phylum           Class                 Order         Family        
# ASV1 "Bacteria" "Proteobacteria" "Gammaproteobacteria" "Coxiellales" "Coxiellaceae"
#        Genus      Species
# ASV1 "Coxiella"     NA   


### DECONTAM  removing contaminants by prevalence, using the blank sample sequenced

sample_data(TickPS)
sample_data(TickPS)$is.neg <- sample_data(TickPS)$Type == "Blank"
sample_data(TickPS)$is.neg
contamdf.prev <- isContaminant(TickPS, method="prevalence", neg="is.neg")
table(contamdf.prev$contaminant)
# FALSE  TRUE 
# 5394    17 

head(which(contamdf.prev$contaminant))
# [1] 205 317 342 377 474 627  --> ID of the cotaminant ASVs

# A more stringent approach:
# threshold=0.5, that will identify as contaminants all sequences thare are more prevalent in 
# negative controls than in positive samples. Lets try using this more aggressive classification threshold rather than the default.

contamdf.prev05 <- isContaminant(TickPS, method="prevalence", neg="is.neg", threshold=0.5)
table(contamdf.prev05$contaminant)

#  FALSE  TRUE 
#  5385    26   --> there were 9 more ASVs that can be deemed as contaminants as they are more prevalent in the Blank than in the samples

TickPS # removing the 26 contaminant taxa retaining only 5385 
TickPS_2 <- prune_taxa(!contamdf.prev05$contaminant, TickPS)
TickPS_2  # 5385 taxa in the otu_table element as expected sans contaminants

# Now we need to remove the ASVs assigned as Eukaryote and NA. Certainly they may contain potential
# eukaryote parasites or could be archaea. Both were not supposed to be amplified by the primer set. We are working with a shorter read seq
# thus affecting the resolution power. 

head(tax_table(TickPS_2)[,1])
TickPS_3<-subset_taxa(TickPS_2, Kingdom=="Bacteria")
TickPS_3  # we retained 2438 ASVs from the 5385 that encompassed eukaryotic and NA ASVs at the Kingdom level

# phyloseq-class experiment-level object
# otu_table()   OTU Table:         [ 2438 taxa and 190 samples ]
# sample_data() Sample Data:       [ 190 samples by 8 sample variables ]
# tax_table()   Taxonomy Table:    [ 2438 taxa by 7 taxonomic ranks ]
# refseq()      DNAStringSet:      [ 2438 reference sequences ]

## Verifying if we have singleton ASVs and samples with just as singleton 

sum(sample_sums(TickPS_3)==0) ## 0 with just a singleton, 0 with zero
sum(taxa_sums(TickPS_3)==0) # 0 ASVs with just one read represented in the table 
# 0 with zero sum
sample_sums(TickPS_3)
head(taxa_sums(TickPS_3))
head(tax_table(TickPS_3))

### Most abundant ASVs in the dataset and their identity
# > head(taxa_sums(TickPS_3))
#       ASV1    ASV2    ASV3    ASV4    ASV6    ASV7 
#     4833348  579194  102740   43111   35868   32117


#         Family                Genus                    Species       
# ASV1 "Coxiellaceae"        "Coxiella"               NA            
# ASV2 "Rickettsiaceae"      "Rickettsia"             "amblyommatis"
# ASV3 "Diplorickettsiaceae" "Rickettsiella"          "grylli"      
# ASV4 "Coxiellaceae"        "Coxiella"               NA            
# ASV6 "Anaplasmataceae"     "Anaplasma"              NA            
# ASV7 "Midichloriaceae"     "Candidatus_Midichloria" NA    


#### Further tax table filtering according to criteria from the previous group publications

# sample prevalence and read abundance filtering thresholds
# "OTU tables were imported into R, and finer filtering was performed. 
# First, we eliminated all OTUs that appeared in no more than two samples. 
# Then, with the reduced data, we further eliminated OTUs with less than 0.5% relative abundance in all ticks."  

## Installing metagMisc to access toolkit of shortcuts for Phyloseq & Ddda2 
install.packages("remotes")
remotes::install_github("vmikk/metagMisc")
devtools::install_github("vmikk/metagMisc")


phyloseq_prevalence_plot(TickPS_3, prev.trh = 0.05, taxcolor = "Phylum",
                         facet = T, point_alpha = 0.7, showplot = T)

# multiple high abundance taxa(>10e+3 reads) are present in aprox 75% of samples
## If we consider 2 samples as a prevalence threshold: 2/189 represents a fraction equal to 0.01

# Removing the Blank from the data set by extracting the subset that belongs to 
# Type = Sample
TickPS_4<-subset_samples(TickPS_3,Type=="Sample")
TickPS_4  ## 189 samples and 2438 taxa 
## Dataset without contaminant ASVs and containing only Bacteria classified sequences

sum(sample_sums(TickPS_4)<1000)
##  0 samples with less than 1000 reads


# Filtering out taxa with less than 10 reads (in total) or not present in at least 1% samples (2/189)
TickPS_5<-phyloseq_filter_prevalence(TickPS_4, prev.trh = 0.01, abund.trh = 10, abund.type = "total", threshold_condition = "OR") 
TickPS_5 # we went from 2438 ASVs to 2224. (retained 91.22%)

# phyloseq-class experiment-level object
# otu_table()   OTU Table:         [ 2224 taxa and 189 samples ]
# sample_data() Sample Data:       [ 189 samples by 8 sample variables ]
# tax_table()   Taxonomy Table:    [ 2224 taxa by 7 taxonomic ranks ]
# refseq()      DNAStringSet:      [ 2224 reference sequences ]


# After filtering we verified once more for singleton ASVs or samples presenting only a singleton

sum(sample_sums(TickPS_5)==1) ## 0 with just a singleton, 0 with zero
sum(taxa_sums(TickPS_5)==1) # 0 ASVs with just one read represented in the table 
# 0 with zero sum

sample_sums(TickPS_5)
head(taxa_sums(TickPS_5))
head(tax_table(TickPS_5))

## Open refseq element to observe its structure
head(refseq(TickPS_5))
#A DNAStringSet instance of length 6
#     width           seq                                        names               
#[1]   150 TACAGAGAGTGCAAGCGTTAATCGG...TAGTAGAGGGAAGTGGAATTTCCGG ASV1
#[2]   150 GACGGGGGGGGCTAGCGTTGTTCGG...TAGTAGGGGATGATGGAATTCCTAG ASV2
#[3]   150 TACAGAGGGTGCAAGCGTTAATCAG...AGATAGAGGGTTGTGGAATTTCCGG ASV3
#[4]   150 TACAGAGAGTGCAAGCGTTAATCGG...TAGTAGAGGGAAGTGGAATTTCCGG ASV4
#[5]   150 TACGGAGGGGGCAAGCGTTGTTCGG...CGGGAGAGGATAGCGGAATTCCTAG ASV6
#[6]   150 TACGAAGGGGGCTAGCGTTACTCGG...CTGTAGGGGGTAGCGGAATTCCTAG ASV7


#################################
# Phylogenetic relatedness is commonly used to inform downstream analyses, especially the calculation of phylogeny-aware distances between microbial communities. 
# The DADA2 sequence inference method is reference-free, so we must construct the phylogenetic tree relating the inferred sequence variants de novo. We begin by 
# performing a multiple-alignment of the inferred sequences.
# load msa package or install it.

if (!requireNamespace("BiocManager", quietly = TRUE))
  install.packages("BiocManager")
BiocManager::install("msa")
a

multT <- msa(refseq(TickPS_5), method="ClustalW", type="dna", order="input")
summary(multT)

# The phangorn package is then used to construct a phylogenetic tree. Here we first construct a neighbor-joining tree, and then fit a GTR+G+I maximum likelihood 
# tree using the neighbor-joining tree as a starting point.

phang.alignT <- as.phyDat(multTalt, type="DNA", names=refseq(TickPS_5))
dmT <- dist.ml(phang.alignT)  # distance measurement
dmT

#NJ tree generation
treeNJT <- NJ(dmT) # Note, tip order == sequence order 
summary(treeNJT)  
plot(treeNJT)

fitT = pml(treeNJT, data=phang.alignT)
## negative edges length changed to 0!


fitGTR_T <- update(fitT, k=4, inv=0.2)
fitGTR_T <- optim.pml(fitGTR_T, model="GTR", optInv=TRUE, optGamma=TRUE,
                      rearrangement = "stochastic", control = pml.control(trace = 0))
detach("package:phangorn", unload=TRUE)

summary(fitGTR_T)
plot(fitGTR_T$tree)
fitGTR_T$data

#####______________________________________
# Creating a new Phyloseq object containing the Phylogeny tree
# Merge TickPS_5 with phylogenetic tree to obtain a new phyloseq objetc used to depict phylogenetic relations combined with abundance patterns 
# and also analyzing Betadiversity with Unifrac dist

TickPS_6 <- merge_phyloseq(TickPS_5alt,phy_tree(fitGTR_T$tree))
TickPS_6

## Create subsets representing the data by Sex for downstream analyses 
sample_data(TickPS_6)

## subsets by Sex

# Male tick subset
TickPS_M<-subset_samples(TickPS_6,Sex == "M")
TickPS_M # 91 samples

sum(sample_sums(TickPS_M)==1) # zero samples with 0 & 1 count
sum(taxa_sums(TickPS_M)==0)  # 14 ASVs have 0 reads and 10 ASVs have just 1 read
### We will prune the dataset remaining only with ASVs that have >10 reads in total as we did for the complete dataset upstream

TickPS_M<-prune_taxa(taxa_sums(TickPS_M)>10,TickPS_M)
sum(sample_sums(TickPS_M)==1) # zero samples with 0 & 1 count
sum(taxa_sums(TickPS_M)==0)# zero taxa with 0 & 1 count
TickPS_M   ###  1835 taxa and 91 samples

# Female tick subset
TickPS_F<-subset_samples(TickPS_6,Sex == "F")
TickPS_F # 98 samples

sum(sample_sums(TickPS_F)==1) # zero samples with 0 & 1 count
sum(taxa_sums(TickPS_F)==0)  # 477 ASVs have 0 reads and 294 ASVs have just 1 read
### We will prune the dataset remaining only with ASVs that have >10 reads in total as above

TickPS_F<-prune_taxa(taxa_sums(TickPS_F)>10,TickPS_F)
sum(sample_sums(TickPS_F)==0) # zero samples with 0 & 1 count
sum(taxa_sums(TickPS_F)==0)# zero taxa with 0 & 1 count
TickPS_F   ###  665 taxa and 98 samples

### Noteworthy to mention the male ticks have a larger richness in terms of unique sequences (ASVs)
## we will see if this traduces to significant alpha div differences when comparing between them and also when segmenting the data set by "region" or evluating this varaible via GLM on alpha div indices

#####################

## Making a data frame from the Phyloseq objects for barplot representations downstream
DF_TickPS_6<-psmelt(TickPS_6)
DF_TickPS_6

# The sample variables: Species have been renamed to "sample_Species"
# to avoid conflicts with taxonomic rank names

DF_TickPS_6_M<-psmelt(TickPS_M)
DF_TickPS_6_M

# The sample variables: Species have been renamed to "sample_Species"
# to avoid conflicts with taxonomic rank names

DF_TickPS_6_F<-psmelt(TickPS_F)
DF_TickPS_6_F

# The sample variables: Species have been renamed to "sample_Species"
# to avoid conflicts with taxonomic rank names

### ALPHA DIVERSITY

alphaTick_full<-estimate_richness(TickPS_6,split=T,measures=c("Observed", "Shannon", "Simpson","Chao1","ACE"))
alphaTick_full
sample_data(TickPS_6)
alphaTick_full$Sex<-sample_data(TickPS_6)$Sex
alphaTick_full$Region<-sample_data(TickPS_6)$Region
alphaTick_full$State<-sample_data(TickPS_6)$State
alphaTick_full$Reads<-sample_sums(TickPS_6)
alphaTick_full$LOGReads<-log(alphaTick_full$Reads,base = exp(10))
write.table(alphaTick_full, "AlphaDiv Ticks.txt", sep="\t", quote=F)
alphaTick_full

#####stats ran with rstatix and ggpubr packages to verify results of both packages

## group data by region and compare between sex, and group by sex and compare between regions
#Simpson
stat.testRvS <- alphaTick_full %>%
  group_by(Region) %>%
  wilcox_test(Simpson ~ Sex) %>%
  adjust_pvalue() %>%
  add_significance("p.adj")
stat.testRvS

stat.testRvS2 <- alphaTick_full %>%
  group_by(Sex) %>%
  wilcox_test(Simpson ~ Region) %>%
  adjust_pvalue() %>%
  add_significance("p.adj")
stat.testRvS2  ### No significant differences between same sex ticks by region for Simpson index

# Observed
stat.testRvSObs <- alphaTick_full %>%
  group_by(Region) %>%
  wilcox_test(Observed ~ Sex) %>%
  adjust_pvalue() %>%
  add_significance("p.adj")
stat.testRvSObs

stat.testRvSObs2 <- alphaTick_full %>%
  group_by(Sex) %>%
  wilcox_test(Observed ~ Region) %>%
  adjust_pvalue() %>%
  add_significance("p.adj")
stat.testRvSObs2 # The only significant difference is detected between female ticks in the HC vs W comparison


### K-W tests of significance
alphaTick_full
#Simpson
compare_means(Simpson ~ Sex,  data = alphaTick_full, method = "kruskal.test",
              group.by = "Region",p.adjust.method="fdr")
# Print table to add as supplementary material for the manuscript

compare_means(Simpson ~ Region,  data = alphaTick_full, method = "kruskal.test",
              group.by = "Sex",p.adjust.method="fdr")  #NS
# Print table to add as supplementary material for the manuscript

#Observed
compare_means(Observed ~ Sex,  data = alphaTick_full, method = "kruskal.test",
              group.by = "Region",p.adjust.method="fdr")

# Print table to add as supplementary material for the manuscript

compare_means(Observed ~ Region,  data = alphaTick_full, method = "kruskal.test",
              group.by = "Sex",p.adjust.method="fdr")  

# Print table to add as supplementary material for the manuscript


# Dunn tests: post-hoc pairwise multiple comparisons...adequate follow up to K-W test
#the Wilcoxon rank sum test, itself a non-parametric analog of the unpaired t-test, is possibly intuitive, 
#but inappropriate as a post hoc pairwise test, because (1) it fails to retain the dependent ranking that 
#produced the Kruskal-Wallis test statistic, and (2) it does not incorporate the pooled variance estimate 
#implied by the null hypothesis of the Kruskal-Wallis test.

alphaTick_full %>%
  group_by(Region) %>%
  dunn_test(Simpson ~ Sex) # Significant differences between different sex ticks when compared for each region


# Boxplot depiction od alpha diversity: Observed ASVs and Simpson index

# variable containing the compared categories to be passed on to stat_compare_means function of ggpubr
my_comparisonstick <- list( c("F", "M"))
my_comparisonstick

# Simpson
ggboxplot(alphaTick_full, x = "Sex", y = "Simpson",
          color = "Sex", palette = "jco", facet.by = "Region", add = "jitter",add.params = list(size = 0.3, jitter = 0.2))+
  stat_compare_means(comparisons = my_comparisonstick, label="p.signif",label.x = 1.5, label.y = 0.8)

## Verify the COLOR choice to make sure it us colorblind friendly

## We are only publishing the Simpson index boxplot to prioritize other figures and present the alpha div data analysis as tables and within the text
# particularly referring to the observed ASVs and applying GLM analysis to the data evaluating the effect of variables such as region and sex


# Barplot depicting the community structure of each sample using facets to distribute the space of the figure by sex and regions 

## Cleanup 
## Data set without the ASVs classified as Bacteria but NA at the Order level. We are removing them for clarity in the barplot though this dataset is not the one used to evaluate
## alpha / beta diversities nor other downstream analyses

TickPS_6_Order_NA <- subset_taxa(TickPS_6, !is.na(Order) & !Order %in% c("", "NA"))
TickPS_6_Order_NA  # now we have 2006 taxa from the 2224 the TickPS_6 dataset contains

# Transforming the dataset to proportions 
propTickPS_6<-transform_sample_counts(TickPS_6_Order_NA, function(x) x / sum(x) )
propTickPS_6 #189 samples and 2006 from the 2224 taxa (218 ASVs are NA's at the Order level) 
sample_sums(propTickPS_6) # all samples sum 1

dataTicks_glomorder <- psmelt(propTickPS_6) # create dataframe from phyloseq object
dataTicks_glomorder$Order <- as.character(dataTicks_glomorder$Order) #convert to character
dataTicks_glomorder$Order
head(dataTicks_glomorder)
tail(dataTicks_glomorder)
write.table(dataTicks_glomorder, "Relative abundances of tick associated microbiota agglomerated at order rank.txt", sep="\t", quote=F)

#simple way to rename genera with < 1% abundance
dataTicks_glomorder$Order[dataTicks_glomorder$Abundance < 0.01] <- "< 1% abund."

pTickOrd <- ggplot(data=dataTicks_glomorder, aes(x=Sample, y=Abundance, fill=Order))

# Color hue created to have contrasting colors for the figure
# Load package "hues"(installed from CRAN)

colorTickhue<-iwanthue(27)  #rather than random, this are optically distinct colors
colorTickhue


# Or use the Iwanthue website to create a colorblind palette
colorhue2<-c("#7863da", "#82c443","#7545b8", "#62ee90", "#9e0079", "#008024", "#d889ff", "#90ab16", "#0180f0", "#ef862f",
"#0173bf", "#df4e39","#01e0e1","#9e0050","#01b076","#4a004a", "#85e8af", "#550039", "#495800",
"#ff75aa","#8e3c00","#8caeff", "#ffb184", "#7b002e", "#ff9489", "#ff87a7", "#ff7a67")


# Barplot by Sex tick mocrobiome with contrasting hue
pTickOrd + geom_bar(aes(), stat="identity", position="stack") +
  scale_fill_manual(values = colorhue2) +
  theme(legend.position="bottom",axis.text.x = element_text(size=0,face="bold")) + guides(fill=guide_legend(nrow=9))+
  facet_grid(~Sex, scales = "free_x" )


# Data set transformations to evaluate beta diversity

#Hellinger to remove weight of low count taxa and many zeroes
Hellinger_TickPS_6full<-TickPS_6
otu_table(Hellinger_TickPS_6full)<-otu_table(decostand(otu_table(Hellinger_TickPS_6full),method= "hellinger"),taxa_are_rows=F)
dim(otu_table(Hellinger_TickPS_6)) 
otu_table(Hellinger_TickPS_6)[1:5,1:5]


# Betadiversity
# Install devtools in order to install and load pairwise.adonis2 packages

install.packages('devtools')
install_github("pmartinezarbizu/pairwiseAdonis/pairwiseAdonis")

# Install MCtools to generate the PCoA plots
devtools::install_github("leffj/mctoolsr")


# Creating a labels variable to be used in permanova, indicator ASV and PCoA analyses
Ticklabels<-sample_data(Hellinger_TickPS_6full)$Sex
Ticklabels<-as.data.frame(Ticklabels)
Ticklabels$Region<-sample_data(Hellinger_TickPS_6full)$Region
Ticklabels
colnames(Ticklabels)<-c("Sex","Region")
Ticklabels

# creating labels for each of the "by sex" phyoseq objects and extra data frames for other analyses
TickPS_F
TickPS_M

TicklabelsF<-dplyr::filter(Ticklabels,Ticklabels$Sex == "F")
TicklabelsF
TicklabelsM<-dplyr::filter(Ticklabels,Ticklabels$Sex == "M")
TicklabelsM

rownames(TicklabelsF)<-rownames(Agglomerated_FAM_Fbinary)
TicklabelsF$sampleID<-rownames(Agglomerated_FAM_Fbinary)
TicklabelsF

rownames(TicklabelsM)<-rownames(Agglomerated_FAM_Mbinary)
TicklabelsM$sampleID<-rownames(Agglomerated_FAM_Mbinary)
TicklabelsM

# We also created binary dataframes to explore other distances such as Jaccard an run co-occurrance tests
TFbinary = as.data.frame(ifelse(otu_table(TickPS_F)>0,1,0))
TFbinary

TMbinary = as.data.frame(ifelse(otu_table(TickPS_M)>0,1,0))
TMbinary

rownames(TicklabelsF)<-rownames(TFbinary)
TicklabelsF$sampleID<-rownames(TFbinary)
TicklabelsF

rownames(TicklabelsM)<-rownames(TMbinary)
TicklabelsM$sampleID<-rownames(TMbinary)
TicklabelsM

# As microbiota data sets are compositional, we are using the Hellinger standardization to generate the proportions reducing the impact of double zeores (joint absences) created by 
# rare ASVs (low in abundance and prevalence) that can be expected on a large data set 

# Female ticks
hellFticks<-TickPS_F
otu_table(hellFticks)[1:5,1:5]
otu_table(hellFticks)<-otu_table(decostand(otu_table(hellFticks),method= "hellinger"),taxa_are_rows=F)
otu_table(hellFticks)[1:5,1:5] ## transformed to reduce weight of rare ASVs (low abundance and prevalence) and double zeroes

dmmcT = calc_dm(t(otu_table(hellFticks)), method = "bray_sq_trans")
dmmcT

TicklabelsF$sampleID
plot_dendrogram(
  dmmcT, metadata_map = TicklabelsF, labels = 'sampleID',
  color_by = 'Region',method="ward.D2"
)
meanRegionBCdistF<-calc_mean_dissimilarities(dmmcT,TicklabelsF,'Region')
meanRegionBCdistF

#       HC        NE         W
#NE 0.4204115                    
#W  0.5315516 0.5605524          
#MW 0.3270336 0.3906553 0.5139614

plot(hclust(meanRegionBCdistF,method = "ward.D2"))

#Male ticks

hellMticks<-TickPS_M
otu_table(hellMticks)[1:5,1:5]
otu_table(hellMticks)<-otu_table(decostand(otu_table(hellMticks),method= "hellinger"),taxa_are_rows=F)
otu_table(hellMticks)[1:5,1:5] ## transformed to reduce weight of rare ASVs (low abundance and prevalence) and double zeroes

dmmcTm = calc_dm(t(otu_table(hellMticks)), method = "bray_sq_trans")
dmmcTm
TicklabelsM
TicklabelsM$sampleID
plot_dendrogram(
  dmmcTm, metadata_map = TicklabelsM, labels = 'sampleID',
  color_by = 'Region',method="ward.D2",main="Males"
)
meanRegionBCdistM<-calc_mean_dissimilarities(dmmcTm,TicklabelsM,'Region')
meanRegionBCdistM

#       HC        NE         W
#NE 0.5159588                    
#W  0.6249241 0.6235558          
#MW 0.4661863 0.4825767 0.5755785


plot(hclust(meanRegionBCdistM,method = "ward.D2"))


# BETADISPER for PCoA ordination 

#Betadisper of BC distance for each of the samples by region (median)
# Females
testBdipserTicksF<-betadisper(dmmcT,TicklabelsF$Region)
testBdipserTicksF
plot(testBdipserTicksF)

#Average distance to median:
#   HC     MW     NE      W 
# 0.2532 0.1780 0.3114 0.4013 

testBdipserTicksM<-betadisper(dmmcTm,TicklabelsM$Region)
testBdipserTicksM
plot(testBdipserTicksM)

# Average distance to median:
#    HC     MW     NE      W 
# 0.3551 0.2368 0.3363 0.4348 

# dispersion tests to address beta diversity / dissimilarity between replicates or samples
# that are characterized as a "group" when using their centroid 

metadispertestFemales <- betadisper(dmmcT,TicklabelsF$Region, type=c("centroid"), bias.adjust = TRUE) 
metadispertestFemales # default is using spatial median as default, but capable of using group centroid too
summary(metadispertestFemales)
scores(metadispertestFemales, display = c("sites", "centroids"), choices=c(1,2))
plot(metadispertestFemales)

metadispertestMales <- betadisper(dmmcTm,TicklabelsM$Region, type=c("centroid"), bias.adjust = TRUE) 
metadispertestMales # default is using spatial median as default, but capable of using group centroid too
summary(metadispertestMales)
scores(metadispertestMales, display = c("sites", "centroids"), choices=c(1,2))
plot(metadispertestMales)


###
# Assessing the significance of the constraints (region grouping)
# Pairwise differences between regions by sex using a function that performs an ANOVA like permutation test

#Females
permutest(metadispertestFemales,pairwise = T,permutations=999)
pstatFem <- permustats(permutest(metadispertestFemales,pairwise = TRUE,permutations=999))
densityplot(pstatFem, scales = list(x = list(relation = "free")))
qqmath(pstatFem, scales = list(relation = "free"))
summary(pstatFem, interval = 0.98)

#Males
permutest(metadispertestMales,pairwise = T,permutations=999)
pstatMa <- permustats(permutest(metadispertestMales,pairwise = TRUE,permutations=999))
pstatMa
densityplot(pstatMa, scales = list(x = list(relation = "free")))
qqmath(pstatMa, scales = list(relation = "free"))
summary(pstatMa, interval = 0.98)

### Improving the plots
#PLot of B diversity (PCoA) dispersion between replicates and experimental condition (Region) centroid

# Female tick PCoA

metadispertestFemales$eig

labspcoaaxisF <- paste("Dimension", 1:97, "(", 
                       round(100*metadispertestFemales$eig / sum(metadispertestFemales$eig), 2), "%)")
labspcoaaxisF #"Dimension 1 ( 33.1 %)"  "Dimension 2 ( 16.0 %)"

plot(metadispertestFemales,label=T ,ellipse=TRUE,conf=0.5,kind = c("ehull"),hull=FALSE,lwd=2,cex=1,cex.axis = 1.5,cex.lab=1.3,col=c("#000000","#9370DB","#FFD700","#FF0000"),
     main="PCoA analysis of female ticks based on their microbial profiles (colored by collection region)",alpha=100, ylab="PCoA2: 16.0%",xlab="PCoA1: 33.1%")

TicklabelsF$Region
legend("bottomright",title="Region",
       c("HC","MW","NE","W"),ncol=1,cex=1,
       fill=c("#000000","#9370DB","#FFD700","#FF0000"))

# Male tick PCoA

metadispertestMales$eig
labspcoaaxisM <- paste("Dimension", 1:90, "(", 
                       round(100*metadispertestMales$eig / sum(metadispertestMales$eig), 2), "%)")
labspcoaaxisM #"Dimension 1 ( 22.5 %)"  "Dimension 2 ( 16.0 %)"

plot(metadispertestMales, ellipse=TRUE,conf=0.5,kind = c("ehull"),hull=FALSE,lwd=2,cex=1,cex.axis = 1.5,cex.lab=1.3,col=c("#000000","#9370DB","#FFD700","#FF0000"),xlim = c(-0.4,0.8),ylim = c(-0.5,0.3),
     main="PCoA analysis of male ticks based on their microbial profiles (colored by collection region)",alpha=100, ylab="PCoA2: 16.0%",xlab="PCoA1: 22.5%")

legend("bottomright",title="Region",ncol=1,
       c("HC","MW","NE","W"),cex=1,
       fill=c("#000000","#9370DB","#FFD700","#FF0000"))

# PERMANOVA to address the significance of the variances when testing for the effect of sex and region interaction

otusTicks<-otu_table(Hellinger_TickPS_6full)
otusTicks<-as.matrix(otusTicks)
otusTicks[1:5,1:5]
dim(otusTicks)

permTicks<- adonis2(otusTicks~Region*Sex,data= Ticklabels, permutations=99999, method="bray", by= "terms")
permTicks


#     adonis2(formula = otusTicks ~ Region * Sex, data = Ticklabels, permutations = 99999, method = "bray", by = "terms")
#             Df SumOfSqs      R2       F Pr(>F)    
#  Region       3    1.389 0.05696  4.7196  1e-05 ***
#  Sex          1    4.922 0.20186 50.1736  1e-05 ***
#  Region:Sex   3    0.316 0.01296  1.0737 0.3533    
#  Residual   181   17.756 0.72821                   
#  Total      188   24.383 1.00000                   
#  ---
#   Signif. codes:  0 ???***??? 0.001 ???**??? 0.01 ???*??? 0.05 ???.??? 0.1 ??? ??? 1


## pairwise adonis by sex ####
hellMticks
hellFticks
TicklabelsF
TicklabelsM

otusTicksF<-otu_table(hellFticks)
dim(otusTicksF)
otusTicksF<-as.matrix(otusTicksF)
otusTicksF[1:5,1:5]
dim(otusTicksF)

pairwiseTickF<-pairwise.adonis2(otusTicksF~Region,data=TicklabelsF, strata=NULL,permutations=999, method ="bray", by="terms",p.adjust.methods="fdr" )
pairwiseTickF


otusTicksM<-otu_table(hellMticks)
dim(otusTicksM)
otusTicksM<-as.matrix(otusTicksM)
otusTicksM[1:5,1:5]
dim(otusTicksM)

pairwiseTickM<-pairwise.adonis2(otusTicksM~Region,data=TicklabelsM, strata=NULL,permutations=999, method ="bray", by="terms",p.adjust.methods="fdr" )
pairwiseTickM


#####Indicator species or ASV analysis

#Tick by Sex and by region : just change the $ character to switch between variables 

# By sex
indicTicks_Sex<-multipatt(as.data.frame(otu_table(Hellinger_TickPS_6full)),Ticklabels$Sex,duleg=F,func="IndVal.g" ,control=how(nperm=999))
indicTicks_Sex$A
summary(indicTicks_Sex)
indictableT<-indicTicks_Sex$sign
indictableT<-as.data.frame(indictableT)
head(indictableT)
colnames(indictableT)

# making a tibble DF that is easier to navigate including the strength of association, pval and Indic Stat 
head(indicTicks_Sex$str[,2])
Indic_ticksDF<-round((indicTicks_Sex$str[,2]),2)
Indic_ticksDF<-as.data.frame(Indic_ticksDF)
head(Indic_ticksDF)
Indic_ticksDF_pval<-as.matrix(indicTicks_Sex$sign[,5])
Indic_ticksDF_stat<-as.matrix(indicTicks_Sex$sign[,4])

Indic_ticksDF$pval<-Indic_ticksDF_pval
Indic_ticksDF$stat<-Indic_ticksDF_stat
Indic_ticksDF$ASV<-rownames(Indic_ticksDF)
colnames(Indic_ticksDF)<-c("strenght of association","pval","Indic_stat","ASV")
head(Indic_ticksDF)


#Subsetting a table based on p values
# https://www.datanovia.com/en/lessons/subset-data-frame-rows-in-r/
#load dplyr and tidyverse
Indic_ticksDF2<-as_tibble(Indic_ticksDF)
Indic_ticksDF2
rownames(Indic_ticksDF2)


Indic_ticksDF3<-Indic_ticksDF2%>%filter(pval <= 0.05)
Indic_ticksDF3
Indic_ticksDF4<-as.data.frame(Indic_ticksDF3)
Indic_ticksDF4 # 219 ASVs 1 for females and 218 for males

## arranging the dataframe in descending order to filter the 
Indic_ticksDF5<-Indic_ticksDF4 %>%
  arrange(desc(Indic_stat)) # we arranged it to make it easier to visualize how we then filter the indicators with a Stat >0.80 
Indic_ticksDF5
Indic_ticksDF6<-Indic_ticksDF5%>%filter(Indic_stat>= 0.80)
Indic_ticksDF6
colnames(Indic_ticksDF6) # verify the col names to add a row pertaining to the only female indicator ASV
Indic_ticksDF7<-Indic_ticksDF6%>% add_row("strenght of association" = 0.00, "pval" = 0.004,"Indic_stat"=0.30,"ASV"="ASV3918")
Indic_ticksDF7
Indic_ticksDF7$Sex<-c(rep("Male",24),rep("Female",1))
Indic_ticksDF7


ASVstoextractT<-Indic_ticksDF7$ASV
ASVstoextractT
ASVstoextractT<-as.vector(ASVstoextractT)
ASVstoextractT # this is the list the ASV numbers for which we will extract abundances and taxa ID from the phyloseq object

TickPS_M # phyloseq objects for each tick sex
TickPS_F

# OTU table element from the phyloseq objects to obtain the total count per ASV within each dataset and estimate 
# the relative abundaces of each ASV in the femal and male datasets

#males
ASVsFullTickM<-otu_table(TickPS_M)
ASVsFullTickM<-as.data.frame(t(ASVsFullTickM))
ASVsFullTickM[1:5,1:5]

CounterIndicASVsM<-rowSums(ASVsFullTickM)
CounterIndicASVsM<-as.data.frame(CounterIndicASVsM)
head(CounterIndicASVsM)
totalreadsM<-colSums(CounterIndicASVsM)
CounterIndicASVsM_REL<-(CounterIndicASVsM/totalreadsM)*100
head(CounterIndicASVsM_REL)
colSums(CounterIndicASVsM_REL) # sums 100
dim(CounterIndicASVsM_REL) #1835 ASVs in the male data set
CounterIndicASVsM_REL
rownames(CounterIndicASVsM_REL)

CounterIndicASVsM_REL2<-CounterIndicASVsM_REL[c("ASV48"  , "ASV74" ,"ASV63"  , "ASV66"   ,"ASV50"  , "ASV45" ,  "ASV49" ,  "ASV59"  , "ASV58"  , "ASV72"  ,
              "ASV61"  , "ASV36"  , "ASV87"  , "ASV60"  , "ASV83" ,  "ASV81"  , "ASV43" ,  "ASV70"  , "ASV73"  , "ASV82"  ,
              "ASV64" ,  "ASV86"  , "ASV30"  , "ASV68","ASV3918"),]
CounterIndicASVsM_REL2
CounterIndicASVsM_REL2<-as.data.frame(CounterIndicASVsM_REL2)
dim(CounterIndicASVsM_REL2) # 24 rows (the indicator ASVs)
colnames(CounterIndicASVsM_REL2)<-c("relative abundance")
rownames(CounterIndicASVsM_REL2)<-c("ASV48"  , "ASV74" ,"ASV63"  , "ASV66"   ,"ASV50"  , "ASV45" ,  "ASV49" ,  "ASV59"  , "ASV58"  , "ASV72"  ,
                                    "ASV61"  , "ASV36"  , "ASV87"  , "ASV60"  , "ASV83" ,  "ASV81"  , "ASV43" ,  "ASV70"  , "ASV73"  , "ASV82"  ,
                                    "ASV64" ,  "ASV86"  , "ASV30"  , "ASV68","ASV3918")

CounterIndicASVsM_REL2[25,1]<-0 ## this ASV is exclusive from the female dataset
CounterIndicASVsM_REL2

#females--> Indicator ASV has a total of 9 reads in the data set so we have to filter the phyloseq object 
# to allow for this ASV to be included. We create a v2 of the female only phyloseq for this purpose 
TickPS_F2<-subset_samples(TickPS_6,Sex == "F")
TickPS_F2 # 98 samples

TickPS_F2<-prune_taxa(taxa_sums(TickPS_F2)>=9,TickPS_F2)
sum(sample_sums(TickPS_F2)==0) # zero samples with 0 & 1 count
sum(taxa_sums(TickPS_F2)==0)# zero taxa with 0 & 1 count
TickPS_F2

ASVsFullTickF<-otu_table(TickPS_F2)
ASVsFullTickF<-as.data.frame(t(ASVsFullTickF))
ASVsFullTickF[1:5,1:5]

CounterIndicASVsF<-rowSums(ASVsFullTickF)
CounterIndicASVsF<-as.data.frame(CounterIndicASVsF)
head(CounterIndicASVsF)
totalreadsF<-colSums(CounterIndicASVsF)
CounterIndicASVsF_REL<-(CounterIndicASVsF/totalreadsF)*100
head(CounterIndicASVsF_REL)
colSums(CounterIndicASVsF_REL) # sums 100
dim(CounterIndicASVsF_REL) #748 ASVs in this dataset
CounterIndicASVsF_REL # indic taxa identified by the multipatt function is present an can be used to depict the relatove abundace of it in a female dataset
rownames(CounterIndicASVsF_REL)

CounterIndicASVsF_REL2<-CounterIndicASVsF_REL[c("ASV48"  , "ASV74" ,"ASV63"  , "ASV66"   ,"ASV50"  , "ASV45" ,  "ASV49" ,  "ASV59"  , "ASV58"  , "ASV72"  ,
                                                "ASV61"  , "ASV36"  , "ASV87"  , "ASV60"  , "ASV83" ,  "ASV81"  , "ASV43" ,  "ASV70"  , "ASV73"  , "ASV82"  ,
                                                "ASV64" ,  "ASV86"  , "ASV30"  , "ASV68","ASV3918"),]

CounterIndicASVsF_REL2
CounterIndicASVsF_REL2<-as.data.frame(CounterIndicASVsF_REL2)
dim(CounterIndicASVsF_REL2) # 24 rows (the indicator ASVs)
colnames(CounterIndicASVsF_REL2)<-c("relative abundance")
rownames(CounterIndicASVsF_REL2)<-c("ASV48"  , "ASV74" ,"ASV63"  , "ASV66"   ,"ASV50"  , "ASV45" ,  "ASV49" ,  "ASV59"  , "ASV58"  , "ASV72"  ,
                                    "ASV61"  , "ASV36"  , "ASV87"  , "ASV60"  , "ASV83" ,  "ASV81"  , "ASV43" ,  "ASV70"  , "ASV73"  , "ASV82"  ,
                                    "ASV64" ,  "ASV86"  , "ASV30"  , "ASV68","ASV3918")

CounterIndicASVsF_REL2


#taxa IDs for the indicator ASVs
TaxIndic<-tax_table(TickPS_6)
TaxIndic<-as.data.frame(TaxIndic)
TaxIndic[1:5,1:7]

TaxIndic2<-TaxIndic[c("ASV48"  , "ASV74" ,"ASV63"  , "ASV66"   ,"ASV50"  , "ASV45" ,  "ASV49" ,  "ASV59"  , "ASV58"  , "ASV72"  ,
                 "ASV61"  , "ASV36"  , "ASV87"  , "ASV60"  , "ASV83" ,  "ASV81"  , "ASV43" ,  "ASV70"  , "ASV73"  , "ASV82"  ,
                 "ASV64" ,  "ASV86"  , "ASV30"  , "ASV68"  , "ASV3918"),]
TaxIndic2
TaxIndic3<-c(rep("p__Proteobacteria",7), "p__Proteobacteria, c__Gammaproteobacteria",rep("p__Proteobacteria",14),"p__Proteobacteria, g__Haemophilus","p__Actinobacteria, g__Rothia","p__Acidobacteria")


Indic_ticksDF7$Relative_abundance_Males<-CounterIndicASVsM_REL2$`relative abundance`
Indic_ticksDF7$Relative_abundance_Females<-CounterIndicASVsF_REL2$`relative abundance`
Indic_ticksDF7

# Dotplot with dotchart function from the hmisc package

Indic_ticksDF7$Sex <- factor(Indic_ticksDF7$Sex) # it must be a factor
Indic_ticksDF8<-Indic_ticksDF7
Indic_ticksDF8$color[Indic_ticksDF8$Sex=='Male'] <- "blue"
Indic_ticksDF8$color[Indic_ticksDF8$Sex=='Female'] <- "orange"
Indic_ticksDF8

dotchart(Indic_ticksDF8$Indic_stat,labels=Indic_ticksDF8$ASV,cex=.7,groups= Indic_ticksDF8$Sex,
         main="Indicator stat ticks by sex",
         xlab="Indicator stat", gcolor="black", color=Indic_ticksDF8$color)


#Adding the taxonomic identity of each ASV
Indic_ticksDF9<-Indic_ticksDF7
Indic_ticksDF9$`Taxonomy ID`<-TaxIndic3    # created another column with a clearer name than TaxID
Indic_ticksDF9
colnames(Indic_ticksDF9)[6]<-c("Relative abundance %")
colnames(Indic_ticksDF9)[3]<-c("Indicator value")
colnames(Indic_ticksDF9)

colorTaxID = c( "#ff8ba3",
                "#00903b",
                "#d6d0ff",
                "#a01caa",
                "#7515cb")


# geom_point plot
Indic_ticksDF9
Indic_ticksDF9 %>%
  mutate(ASV = fct_reorder(ASV,`Relative abundance %`)) %>%
ggplot( aes(x = `Indicator value`, y = ASV))+geom_point(aes(size = `Relative abundance %`, fill = `Taxonomy ID`,stroke=1), alpha = 1,shape=21)+
  scale_size_continuous(limits = c(0.0000, 1), range = c(4,13), breaks = c(0.0001,0.10,0.20))+ guides(fill = guide_legend(override.aes = list(size=6)))+
  scale_fill_manual(values = colorTaxID)+theme_minimal()+
  theme(panel.grid = element_line(color = "grey",
                                  size = 0.3,
                                  linetype = 1),
        panel.grid.minor.x =element_blank(),panel.grid.major.x =element_blank())+
        theme(legend.text = element_text(size = 10),
        axis.text.x = element_text(colour = "black", size = 10,vjust = 0.3, hjust = 1),
        axis.text.y = element_text(colour = "black" , size = 10))
  
#test with shape by Sex category and ordered by indicator value
Indic_ticksDF9 %>%
  mutate(ASV = fct_reorder(ASV,`Indicator value`)) %>%
  ggplot( aes(x = `Indicator value`, y = ASV))+geom_point(aes(size = `Relative abundance %`, colour = `Taxonomy ID`,shape=`Sex`,stroke=1), alpha = 1)+
  scale_size_continuous(limits = c(0.0000, 1), range = c(4,12), breaks = c(0.0001,0.10,0.20))+ guides(colour = guide_legend(override.aes = list(size=6)))+
  guides(shape = guide_legend(override.aes = list(size=6)))+scale_color_manual(values = colorTaxID)+theme_minimal()+
  theme(panel.grid = element_line(color = "grey",
                                  size = 0.3,
                                  linetype = 1),
        panel.grid.minor.x =element_blank(),panel.grid.major.x =element_blank())+
  theme(legend.text = element_text(size = 10),
        axis.text.x = element_text(colour = "black", size = 10,vjust = 0.3, hjust = 1),
        axis.text.y = element_text(colour = "black" , size = 10))


####
### Indicator ASVs per region
indicTicks_Region<-multipatt(as.data.frame(otu_table(Hellinger_TickPS_6full)),Ticklabels$Region,duleg=T,func="IndVal.g" ,control=how(nperm=999))
summary(indicTicks_Region)
# NO indicator ASV for the HC region

indicTicks_Region$sign
indictableTreg<-indicTicks_Region$sign
indictableTreg<-as.data.frame(indictableTreg)
head(indictableTreg)
# making a tibble DF that is easier to navigate including the pval and Indic Stat 
indicTicks_Region$str
Indic_ticksDFreg<-round((indicTicks_Region$str[,]),2)
Indic_ticksDFreg<-as.data.frame(Indic_ticksDFreg)
head(Indic_ticksDFreg)

Indic_ticksDFreg_pval<-as.matrix(indicTicks_Region$sign[,7])
Indic_ticksDFreg_stat<-as.matrix(indicTicks_Region$sign[,6])

Indic_ticksDFreg$pval<-Indic_ticksDFreg_pval
Indic_ticksDFreg$stat<-Indic_ticksDFreg_stat
Indic_ticksDFreg$ASV<-rownames(Indic_ticksDFreg)
colnames(Indic_ticksDFreg)[5:7]
colnames(Indic_ticksDFreg)[5:7]<-c("pval","Indic_stat","ASV")
head(Indic_ticksDFreg)


#Subsetting a table based on p values
# https://www.datanovia.com/en/lessons/subset-data-frame-rows-in-r/
#load dplyr and tidyverse
Indic_ticksDF2reg<-as_tibble(Indic_ticksDFreg)
Indic_ticksDF2reg
rownames(Indic_ticksDF2reg)


Indic_ticksDF3reg<-Indic_ticksDF2reg%>%filter(pval <= 0.05)
Indic_ticksDF3reg
Indic_ticksDF4reg<-as.data.frame(Indic_ticksDF3reg)
head(Indic_ticksDF4reg) # 

## arranging the dataframe in descending order to filter the ASVs based on an arbitrary indic stat threshold
Indic_ticksDF5reg<-Indic_ticksDF4reg %>%
  arrange(desc(Indic_stat)) # we arranged it to make it easier to visualize how we then filter the indicators with a Stat >0.80 
head(Indic_ticksDF5reg)
Indic_ticksDF6reg<-Indic_ticksDF5reg%>%filter(Indic_stat>= 0.6)
dim(Indic_ticksDF6reg) #this cutoff filters the DF to 38 indicators which we need to associate to each region afterwards from the multipatt summary above 
                       # we can report the number of significant indicators per region in text.

summary(indicTicks_Region)
Indic_ticksDF6reg
#MW ASV32
#NE ASV2 
#W the rest
Indic_ticksDF6reg$Region<-c(rep("West",22),"Northeast",rep("West",9),"Midwest",rep("West",5))
Indic_ticksDF6reg

# shape assigned per region for the plot
Indic_ticksDF6reg$shape[Indic_ticksDF6reg$Region=='West'] <- 22
Indic_ticksDF6reg$shape[Indic_ticksDF6reg$Region=='Northeast'] <- 21
Indic_ticksDF6reg$shape[Indic_ticksDF6reg$Region=='Midwest'] <- 24
Indic_ticksDF6reg

# ASVs to extract their relative abundances from the full dataset
ASVstoextractTW<-Indic_ticksDF6reg$ASV
ASVstoextractTW
ASVstoextractTW<-as.vector(ASVstoextractTW)
ASVstoextractTW # this is the list the ASV numbers for which we will extract abundances and taxa ID from the phyloseq object containing data from Western ticks

ASVstoextractTNE<-c("ASV2")
ASVstoextractTMW<-c("ASV32")

# Subset phyloseq for each region in order to extract relative abundances of the indicator ASVs
HCTicks<-subset_samples(TickPS_6,Region=="HC")
WTicks<-subset_samples(TickPS_6,Region=="W")
NETicks<-subset_samples(TickPS_6,Region=="NE")
MWTicks<-subset_samples(TickPS_6,Region=="MW")

# OTU table element from the phyloseq objects to obtain the total count per ASV within each dataset and estimate 
# the relative abundaces of each ASV in the femal and male datasets

#West
ASVscountTW<-otu_table(WTicks)
ASVscountTW<-as.data.frame(t(ASVscountTW))
ASVscountTW[1:5,1:5]

CounterIndicASVsTW<-rowSums(ASVscountTW)
CounterIndicASVsTW<-as.data.frame(CounterIndicASVsTW)
head(CounterIndicASVsTW)
totalreadsTW<-colSums(CounterIndicASVsTW)
CounterIndicASVsTW_REL<-(CounterIndicASVsTW/totalreadsTW)*100
head(CounterIndicASVsTW_REL)
colSums(CounterIndicASVsTW_REL) # sums 100
dim(CounterIndicASVsTW_REL) 
CounterIndicASVsTW_REL
rownames(CounterIndicASVsTW_REL)

ASVstoextractTW
CounterIndicASVsTW_REL2<-CounterIndicASVsTW_REL[c(ASVstoextractTW),]
CounterIndicASVsTW_REL2
CounterIndicASVsTW_REL2<-as.data.frame(CounterIndicASVsTW_REL2)
dim(CounterIndicASVsTW_REL2) # 
colnames(CounterIndicASVsTW_REL2)<-c("relative abundance")
rownames(CounterIndicASVsTW_REL2)<-c(ASVstoextractTW)
CounterIndicASVsTW_REL2

#NE
ASVscountTNE<-otu_table(NETicks)
ASVscountTNE<-as.data.frame(t(ASVscountTNE))
ASVscountTNE[1:5,1:5]

CounterIndicASVsTNE<-rowSums(ASVscountTNE)
CounterIndicASVsTNE<-as.data.frame(CounterIndicASVsTNE)
head(CounterIndicASVsTNE)
totalreadsTNE<-colSums(CounterIndicASVsTNE)
CounterIndicASVsTNE_REL<-(CounterIndicASVsTNE/totalreadsTNE)*100
head(CounterIndicASVsTNE_REL)
colSums(CounterIndicASVsTNE_REL) # sums 100
dim(CounterIndicASVsTNE_REL) 

ASVstoextractTNE
CounterIndicASVsTNE_REL2<-CounterIndicASVsTNE_REL[c(ASVstoextractTNE),]
CounterIndicASVsTNE_REL2
CounterIndicASVsTNE_REL2<-as.data.frame(CounterIndicASVsTNE_REL2)
dim(CounterIndicASVsTNE_REL2) # 
colnames(CounterIndicASVsTNE_REL2)<-c("relative abundance")
rownames(CounterIndicASVsTNE_REL2)<-c(ASVstoextractTNE)
CounterIndicASVsTNE_REL2

#Midwest
ASVscountTMW<-otu_table(MWTicks)
ASVscountTMW<-as.data.frame(t(ASVscountTMW))
ASVscountTMW[1:5,1:5]

CounterIndicASVsTMW<-rowSums(ASVscountTMW)
CounterIndicASVsTMW<-as.data.frame(CounterIndicASVsTMW)
head(CounterIndicASVsTMW)
totalreadsTMW<-colSums(CounterIndicASVsTMW)
CounterIndicASVsTMW_REL<-(CounterIndicASVsTMW/totalreadsTMW)*100
head(CounterIndicASVsTMW_REL)
colSums(CounterIndicASVsTMW_REL) # sums 100
dim(CounterIndicASVsTMW_REL) 

ASVstoextractTMW
CounterIndicASVsTMW_REL2<-CounterIndicASVsTNE_REL[c(ASVstoextractTMW),]
CounterIndicASVsTMW_REL2
CounterIndicASVsTMW_REL2<-as.data.frame(CounterIndicASVsTMW_REL2)
dim(CounterIndicASVsTMW_REL2) # 
colnames(CounterIndicASVsTMW_REL2)<-c("relative abundance")
rownames(CounterIndicASVsTMW_REL2)<-c(ASVstoextractTMW)
CounterIndicASVsTMW_REL2

# inserting the relative abundances for the MW and NE indicators into the DF we wil use to plot the data
rownames(CounterIndicASVsTW_REL2)
# inserting NE indicator
CounterIndicASVsTNE_REL2
CounterIndicASVsTW_REL2[23,1]<-12.23858 
CounterIndicASVsTW_REL2

## inserting MW indicator
CounterIndicASVsTMW_REL2
CounterIndicASVsTW_REL2[33,1]<-0.1030011 
CounterIndicASVsTW_REL2

taxIDregs<-rownames(CounterIndicASVsTW_REL2)
taxIDregs

#Ploting the indicators per region with their relative abundance as size marker representing their abundance in each region

#taxa IDs for the indicator ASVs
TaxIndic<-tax_table(TickPS_6)
TaxIndic<-as.data.frame(TaxIndic)
TaxIndic[1:5,1:7]

TaxIndicReg<-TaxIndic[c(taxIDregs),]
TaxIndicReg
TaxIndicRegFam<-TaxIndicReg$Family
TaxIndicRegFam
TaxIndicReg2<-c(rep("p__Proteobacteria",2),"p__Bacteroidetes",rep("p__Proteobacteria",2),"p__Bacteroidetes","p__Acidobacteria",rep("p__Proteobacteria",6),"p__Bacteroidetes",rep("p__Proteobacteria",6),"p__Bacteroidetes",rep("p__Proteobacteria",4),"p__Bacteroidetes","p__Proteobacteria",rep("p__Bacteroidetes",2),rep("p__Proteobacteria",2),"p__Verrucomicrobiota",rep("p__Proteobacteria",6))
TaxIndicReg2
# load stringr
taxregcomb<- str_c(TaxIndicReg2,", f__",TaxIndicRegFam)
taxregcomb
taxregcomb[24]<-c("p__Proteobacteria, o__Rickettsiales")
taxregcomb[33]<-c("p__Proteobacteria, o__Rickettsiales")
taxregcomb

# Dotplot with dotchart function from the hmisc package
Indic_ticksDF6reg
Indic_ticksDF6reg$Region <- factor(Indic_ticksDF6reg$Region) # it must be a factor
Indic_ticksDF7reg<-Indic_ticksDF6reg
Indic_ticksDF7reg$color[Indic_ticksDF7reg$Region=='West'] <- "#FF0000"
Indic_ticksDF7reg$color[Indic_ticksDF7reg$Region=='Northeast'] <- "#FFD700"
Indic_ticksDF7reg$color[Indic_ticksDF7reg$Region=='Midwest'] <- "#9370DB"
Indic_ticksDF7reg$color

#dotchart
dotchart(Indic_ticksDF7reg$Indic_stat,labels=Indic_ticksDF7reg$ASV,cex=.7,groups= Indic_ticksDF7reg$Region,
         main="Indicator stat ticks by region",
         xlab="Indicator stat", gcolor="black", color=Indic_ticksDF7reg$color)


#Adding the taxonomic identity of each ASV
Indic_ticksDF8reg<-Indic_ticksDF6reg
Indic_ticksDF8reg$`Taxonomy ID`<-taxregcomb    
Indic_ticksDF8reg
# relative abundances columns
CounterIndicASVsTW_REL2
Indic_ticksDF8reg$`Relative abundance %`<-CounterIndicASVsTW_REL2$`relative abundance`
Indic_ticksDF8reg
colnames(Indic_ticksDF8reg)[6]<-c("Indicator value")
colnames(Indic_ticksDF8reg)

colorTaxIDreg = c( "#008fff",
                "#d6d0ff",
                "#a01caa",
                "#7515cb","#ff9c80",
                "#714a34",
                "#fdc8c5",
                "#90342e",
                "#c39b82",
                "#c4646a",
                "#ae6732",
                "#b47476", "#00a540")


# geom_point plot
Indic_ticksDF8reg %>%
  mutate(ASV = fct_reorder(ASV,`Indicator value`)) %>%
  ggplot( aes(x = `Indicator value`, y = ASV))+geom_point(aes(colour = `Taxonomy ID`, shape = `Region`,stroke=1),size = 4.1)+
  guides(colour = guide_legend(override.aes = list(size=6)))+guides(Region = guide_legend(override.aes = list(size=6)))+
  theme_minimal()+scale_color_manual(values = colorTaxIDreg)+
  theme(panel.grid = element_line(color = "grey",
                                  size = 0.3,
                                  linetype = 1),
  panel.grid.minor.x =element_blank(),panel.grid.major.x =element_blank())+
  theme(legend.text = element_text(size = 10),
        axis.text.x = element_text(colour = "black", size = 10,vjust = 0.3, hjust = 1),
        axis.text.y = element_text(colour = "black" , size = 9))

#scale_size_continuous(limits = c(0.00,12), range = c(0,12), breaks = c(0.001,0.2,12))+


## " A statistic commonly used to determine the association (also known as fidelity, 
#not to be confounded with the indicator value component B) between species and niche types 
# is Pearson???s phi coefficient of association [Chytr?? et al., 2002]. 
# This coefficient is a measure of the correlation between two binary vectors. It is possible to calculate the 
# phi coefficient in multipatt after transforming our community data to presence-absence format".


## Point biserial correlation coefficient
Binaryticks = ifelse(as.data.frame((otu_table(Hellinger_TickPS_6full)))>0,1,0)
Binaryticks[1:5,1:5]
phi_ticks_sex = multipatt(Binaryticks,Ticklabels$Sex, func = "r.g",control = how(nperm=999),duleg=T)
round(phi_ticks_sex$str[,2],3)
summary(phi_ticks_sex)


# making a tibble DF that is easier to navigate including the strength of association, pval and Indic Stat 
#males
evasion_ticks<-round((phi_ticks_sex$str[,2]),2)
evasion_ticks<-as.data.frame(evasion_ticks)
head(evasion_ticks)
evasion_ticks_pval<-as.matrix(phi_ticks_sex$sign[,5])
evasion_ticks_stat<-as.matrix(phi_ticks_sex$sign[,4])

#females
evasion_ticksF<-round((phi_ticks_sex$str[,1]),2)
evasion_ticksF<-as.data.frame(evasion_ticksF)
head(evasion_ticksF)
evasion_ticks_pvalF<-as.matrix(phi_ticks_sex$sign[,5])
evasion_ticks_statF<-as.matrix(phi_ticks_sex$sign[,4])


evasion_ticks$pval<-evasion_ticks_pval
evasion_ticks$stat<-evasion_ticks_stat
evasion_ticks$ASV<-rownames(evasion_ticks)
colnames(evasion_ticks)<-c("strenght of association","pval","Indic_stat","ASV")
head(evasion_ticks)


evasion_ticksF$pval<-evasion_ticks_pvalF
evasion_ticksF$stat<-evasion_ticks_statF
evasion_ticksF$ASV<-rownames(evasion_ticksF)
colnames(evasion_ticksF)<-c("strenght of association","pval","Indic_stat","ASV")
head(evasion_ticksF)


#Subsetting a table based on p values
# https://www.datanovia.com/en/lessons/subset-data-frame-rows-in-r/
#load dplyr and tidyverse
evasion_ticks2sex<-as_tibble(evasion_ticks)
evasion_ticks2sex
rownames(evasion_ticks2sex)

evasion_ticks2sexF<-as_tibble(evasion_ticksF)
evasion_ticks2sexF
rownames(evasion_ticks2sexF)


evasion_ticks3sex<-evasion_ticks2sex%>%filter(pval <= 0.05)
evasion_ticks3sex<-evasion_ticks3sex%>%filter(`strenght of association` < 0)
evasion_ticks3sex
evasion_ticks4sex<-as.data.frame(evasion_ticks3sex)
evasion_ticks4sex
class(evasion_ticks4sex)
## 6 ASVs from male ticks with significant negative strength of association regarding female samples

evasion_ticks3sexF<-evasion_ticks2sexF%>%filter(pval <= 0.05)
evasion_ticks3sexF<-evasion_ticks3sexF%>%filter(`strenght of association` < 0)
evasion_ticks3sexF
evasion_ticks4sexF<-as.data.frame(evasion_ticks3sexF)
evasion_ticks4sexF
## 181 ASVs from female ticks with significative negative strenght of association regarding male samples.


### Microeco package to do: Estimation of ecological processes' influence on variance, and Functional profiles per tick sex
# source: https://chiliubio.github.io/microeco/

# Install file2meco to transform phyloseq object to microeco table
devtools::install_github("ChiLiubio/file2meco")

# transform to microeco microtables and tidy the data as suggested in the tutorial to unify taxonomic prefixes and cleanup NA's, unidentified and unknown calls that can 
# influence taxonomic abundance calculations downtream. 

# full dataset
microfull<-phyloseq2meco(TickPS_6)
tidy_taxonomy(microfull$tax_table)
microfull$tidy_dataset()
microfull$tax_table[29,]

# Female tick data
microTSfem<-phyloseq2meco(TickPS_F)
tidy_taxonomy(microTSfem$tax_table)
microTSfem$tidy_dataset()
# Male tick data
microTSmale<-phyloseq2meco(TickPS_M)
tidy_taxonomy(microTSmale$tax_table)
microTSmale$tidy_dataset()

# Tidy dataset to make the OTU and sample information consistent across all files in the dataset object

# Taxa abundance

#males
microTSmale$cal_abund()
#females
microTSfem$cal_abund()


## Using Microeco package to generate DF with beta div measures
# package GUnifrac needed but only works with rooted phylogenetic trees
# thus we extracted the wunifrac dist matrix from the phyloseq objetc.
microfull
microfull$cal_betadiv(unifrac = TRUE)

BrayBdivmatrix<-microfull$beta_diversity$bray
BTNRIBdivmatrix<-microfull$beta_diversity$betaNRI
BTNRIBdivmatrix
JaccardBdivmatrix<-microfull$beta_diversity$jaccard
#wunifrac
Fulltick_wunifrac <- phyloseq::distance(TickPS_6_prop, method = "wunifrac")
Fulltick_wunifrac

### Prospective genomics with full dataset comparing among samples 

FuncTfullIND <- trans_func$new(microfull)
FuncTfullIND
FuncTfullIND$cal_spe_func()
FuncTfullIND$res_spe_func[1:10,50:60]
FuncTfullIND$cal_spe_func_perc(abundance_weighted = F)
FuncTfullIND$res_spe_func_perc
FuncTfullIND$plot_spe_func_perc()

matfunTicksFull<-as.matrix(FuncTfullIND$res_spe_func_perc)
dim(matfunTicksFull)
rownames(matfunTicksFull)

FuncTfullIND$sample_table$Sex
matfunTicksFull2<-t(matfunTicksFull)
matfunTicksFull2

colnames(FuncTfullIND$sample_table)
FuncTfullIND$sample_table[,c(2,5)]
Sideinfo_sex_region<-FuncTfullIND$sample_table[,c(2,5)]
Sideinfo_sex_region
Sideinfo_sex_region<-as.matrix(Sideinfo_sex_region)
Sideinfo_sex_region<-as.data.frame(Sideinfo_sex_region)
Sideinfo_sex_region

ann_colorsFUNCTICK=list(Sex=c('F'="#C0C0C0",'M'="#808000"),Region=c('HC'="#000000",'W'="#FF0000",'NE'="#FFD700",'MW'="#9370DB"))
ann_colorsFUNCTICK


pheatmap(matfunTicksFull2,cluster_rows = T,cluster_cols = T,fontsize_col = 5,fontsize_row=5,annotation_col = Sideinfo_sex_region,annotation_colors =ann_colorsFUNCTICK ,cellwidth = 5,cellheight = 6,clustering_distance_cols = "minkowski",clustering_method = "median",
         color=colorRampPalette(c("white","orange","salmon","red","firebrick")) (1024)) 

# euclidean and median good
# euclidean and single good
# minkowski median is currently the best

### Adding a state color annotation may help as we can then remove sample names
FuncTfullIND$sample_table$State
## 17 states

FuncTfullIND$sample_table[,c(2,4,5)]
Sideinfo_sex_region2<-FuncTfullIND$sample_table[,c(2,4,5)]
Sideinfo_sex_region2
Sideinfo_sex_region2<-as.matrix(Sideinfo_sex_region2)
Sideinfo_sex_region2<-as.data.frame(Sideinfo_sex_region2)
Sideinfo_sex_region2


ann_colorsFUNCTICK2=list(Sex=c('F'="#00BFFF",'M'="#32CD32"),Region=c('HC'="#000000",'W'="#FF0000",'NE'="#FFD700",'MW'="#9370DB"),
                         State=c('AL'="#696969",'AR'="#808080",'FL'="#A9A9A9",'GA'="#D3D3D3",'IN'="#DCDCDC",'KS'="#FF6347",'KY'="#F5F5F5",'MD'="#FFFF00",'MO'="#708090",'MS'="#778899",'NC'="#B0C4DE",'NJ'="#E6E6FA",'OH'="#6A5ACD",'OK'="#B22222",'RI'="#F0E68C",'TN'="#808080",'VA'="#DAA520")                  )
ann_colorsFUNCTICK2


pheatmap(matfunTicksFull2,cluster_rows = T,cluster_cols = T,fontsize_col = 3,fontsize_row=9,annotation_col = Sideinfo_sex_region2,annotation_colors =ann_colorsFUNCTICK2 ,cellwidth = 5,cellheight = 7,clustering_distance_cols = "minkowski",clustering_method = "median",
         color=colorRampPalette(c("white","plum","#9370DB","#483D8B","#4B0082")) (1024)) 


# Identifying functional categories significantly different between tick sex

TFUNCstat.test2 <- compare_means(
  value ~ Sex, data = matfunTicksFull2DF2, group.by = "variable",
  method = "wilcox.test", p.adjust.method = "fdr")
TFUNCstat.test2

TSigCat2<-TFUNCstat.test2 %>% filter(TFUNCstat.test2$p.adj<0.05)
TSigCat2


#### Other beta div indices and analyses:
## Ecological processes driving the variance among microbial communities

# generate trans_nullmodel object
# we only use high abundance OTU with mean relative abundance > 0.0005
BetaFem<-trans_nullmodel$new(microTSfem, filter_thres = 0.0005)
BetaMale<-trans_nullmodel$new(microTSmale, filter_thres = 0.0005)


BetaFem$cal_ses_betampd(runs=500, abundance.weighted = TRUE)
BetaFem$res_ses_betampd

BetaMale$cal_ses_betampd(runs=500, abundance.weighted = TRUE)
BetaMale$res_ses_betampd

# add betaNRI matrix to beta_diversity results

microTSfem$beta_diversity[["betaNRI"]] <- BetaFem$res_ses_betampd
microTSfem$beta_diversity$betaNRI
microTSmale$beta_diversity[["betaNRI"]] <- BetaMale$res_ses_betampd
microTSmale$beta_diversity$betaNRI

#females
BetaFemtransnull_reg <- trans_beta$new(dataset = microTSfem, group = "Region", measure = "betaNRI")
#males
BetaMtransnull_reg <- trans_beta$new(dataset = microTSmale, group = "Region", measure = "betaNRI")


## transform the distance for each group
BetaFemtransnull_reg$cal_group_distance()
BetaMtransnull_reg$cal_group_distance()


FEMbetaplot1reg <- BetaFemtransnull_reg$plot_group_distance(distance_pair_stat = TRUE)
FEMbetaplot1reg + geom_hline(yintercept = -2, linetype = 2) + geom_hline(yintercept = 2, linetype = 2)

Mbetaplot1reg <- BetaMtransnull_reg$plot_group_distance(distance_pair_stat = TRUE)
Mbetaplot1reg + geom_hline(yintercept = -2, linetype = 2) + geom_hline(yintercept = 2, linetype = 2)


#BetaNTI(ses.betamntd) can be used to indicate the phylogenetic terminal turnover (Stegen et al. 2013).
BetaFem$cal_ses_betamntd(runs=500, abundance.weighted = TRUE)
BetaFem$cal_rcbray(runs = 1000)
BetaFem$res_rcbray
BetaFem$res_ses_betamntd

BetaMale$cal_ses_betamntd(runs=500, abundance.weighted = TRUE)
BetaMale$cal_rcbray(runs = 1000)
BetaMale$res_rcbray
BetaMale$res_ses_betamntd


# use betaNTI and rcbray to determine the percentage that can be assigned to ecological processes 
BetaFem$cal_process(use_betamntd = T)
BetaFem$res_process

BetaMale$cal_process(use_betamntd = T)
BetaMale$res_process


###ONLY FEMALES
#> BetaFem$res_process
#            process      percentage
#1    variable selection   0.000000
#2 homogeneous selection   0.610141
#3  dispersal limitation  11.277088
#4 homogeneous dispersal   0.000000
#5                 drift  88.112771


###ONLY MALES
#> BetaMale$res_process
#           process       percentage
#1    variable selection  0.0000000
#2 homogeneous selection  0.0976801
#3  dispersal limitation 12.9670330
#4 homogeneous dispersal  1.0256410
#5                 drift 85.9096459


# assembly processes matrix for plot
femprocess<-as.data.frame(BetaFem$res_process)
maleprocess<-as.data.frame(BetaMale$res_process)

# One fast way of creating a DF for the plot
processmatrix<-femprocess
processmatrix<-processmatrix %>%
  mutate(maleprocess$percentage)
colnames(processmatrix)<-c("Process","Females","Males")
row.names(processmatrix)<-processmatrix[,1]
row.names(processmatrix)
processmatrix<-processmatrix[,2:3]
rownames(processmatrix)
head(processmatrix)
#Bubble plot (the function is no longer available)
bubble_plot(processmatrix,transp=0.9,circ = 25, font.label =(size=15))


# Another easy way of creating a small data frame for figure generation with a bubble plot that allows us to remove the 0 values and manage the size of the elements.

TProcess=c("variable selection", "homogeneus selection","dispersal limitation" ,"homogeneous dispersal" ,"drift" )
TFemalesp=c(0.00,0.61,11.28,0.00,88.11) 
TMalesp=c(0.00,0.10,12.97,1.03,85.90)
EcoProcTicks=data.frame(TProcess,TFemalesp,TMalesp)
colnames(EcoProcTicks)<-c("Process","Females","Males")
EcoProcTicks
EcoProcTicks2<-melt(EcoProcTicks,id = c("Process")) # melt is a function from the reshape package
EcoProcTicks2$variable<-factor(EcoProcTicks2$variable,levels=unique(EcoProcTicks2$variable))
EcoProcTicks2
coloursTbub = c( "#FFA500","#008080")

ggplot(EcoProcTicks2, aes(x = variable, y = Process)) + 
  geom_point(aes(size = value, fill = variable), alpha = 0.55, shape = 21) + 
  scale_size_continuous(limits = c(0.000001, 100), range = c(1,45), breaks = c(1,5,10,80)) + 
  labs( x= "", y = "", size = "Relative Abundance (%)", fill = "")  + 
  theme(legend.key=element_blank(), 
        axis.text.x = element_text(colour = "black", size = 15, face = "bold", angle = 90, vjust = 0.3, hjust = 1), 
        axis.text.y = element_text(colour = "black", face = "bold", size = 15), 
        legend.text = element_text(size = 10, face ="bold", colour ="black"), 
        legend.title = element_text(size = 12, face = "bold"), 
        panel.background = element_blank(), panel.border = element_rect(colour = "black", fill = NA, size = 1.2), 
        legend.position = "right") +  
  scale_fill_manual(values = coloursTbub, guide = FALSE) + 
  scale_y_discrete(limits = rev(levels(EcoProcTicks2$Process)))


           
### Spatial autocorrelation analysis. Using the sample collection coordinates and the beta diversity to evaluate if 
# geographic structuring of the microbiota is correlated to its beta diversity
## Limma package needed

## Obtaining the coordinates for each sample from the metadata in the phyloseq object
BiocManager::install("limma")

colnames(sample_data(TickPS_6))
coordinates<-sample_data(TickPS_6)[,6]
coordinates<-as.matrix(coordinates)
coordinates
coordinates2<-strsplit2(coordinates[,],",")
row.names(coordinates2)<-row.names(coordinates)
coordinates2
class(coordinates2)

# by sex coordinates
coordinatesF<-as.data.frame(coordinates2)
coordinatesF

ticksexID<-alphaTick_2$Sex

coordinatesF$Sex<-alphaTick_2$Sex
coordinatesF

coordinatesM<-filter(coordinatesF, Sex == "M") 
coordinatesM
coordinatesF2<-filter(coordinatesF, Sex == "F")
coordinatesF2

coordinatesM2<-as.matrix(coordinatesM)
coordinatesM2<-coordinatesM2[,1:2]
coordinatesM2

coordinatesF3<-as.matrix(coordinatesF2)
coordinatesF3<-coordinatesF3[,1:2]
coordinatesF3

# Geodist 
#Convert  one  or  two  rectangular  objects  containing  lon-lat  coordinates  into  vector  or  matrix  of geodesic distances in meters.
coordinates2
spatialdist<-geodist (coordinates2, sequential = F, pad = F,measure="vincenty")
spatialdist
dim(spatialdist)
row.names(spatialdist)<-row.names(coordinates2)
colnames(spatialdist)<-row.names(coordinates2)
spatialdist[1:20,1:5]

# writing table of spatial distances that can be added as a supplementary file in case other approaches for correlograms or semivariograms can be used
write.table(spatialdist, "Spatial distances_Vincenty_mts.txt", sep="\t", quote=F)

## mantel correlogram by sex
hellFticks
hellMticks
mantelhellF<-as.matrix(otu_table(hellFticks))
mantelhellM<-as.matrix(otu_table(hellMticks))

# geospatial coordinates used to create the distance classes in the correlogram
dfcoordF<-as.data.frame(coordinatesF3)
head(dfcoordF)

dfcoordM<-as.data.frame(coordinatesM2)
head(dfcoordM)

# vegdist Bray on hellinger abundance matrix
tick.hel.DM2 <- vegdist(mantelhellM,method = "bray")
tick.hel.DF2 <- vegdist(mantelhellF,method = "bray")

# Compute Mantel correlogram with cutoff, Spearman statistic
tick.correlogF <- mantel.correlog(tick.hel.DF2, XY=dfcoordF, nperm=99,r.type="spearman",cutoff=FALSE,mult="fdr")
summary(tick.correlogF)
tick.correlogF   
plot(tick.correlogF)


tick.correlogM <- mantel.correlog(tick.hel.DM2, XY=dfcoordM, nperm=999,r.type="spearman",cutoff=FALSE,mult="fdr")
summary(tick.correlogM)
tick.correlogM
plot(tick.correlogM)

# improving the mantel correlogram plot

DFTF_mantel<-tick.correlogF$mantel.res   
DFTF_mantel<-as.data.frame(DFTF_mantel)
class(DFTF_mantel)
tick.correlogF # determine the alpha values depending on the FDR corrected p values
DFTF_mantel$alpha<-c(1,0.1,0.85,0.85,0.1,0.1,0.85,0.1,0.85,0.1,0.85,0.1,0.1,0.1)
DFTF_mantel


  MantCorF<-DFTF_mantel %>%
  tail(100) %>%
ggplot( aes(x=class.index, y=Mantel.cor)) +
  geom_line( color="grey") +
  geom_point(shape=21, color="black", fill="black", size=6, alpha=DFTF_mantel$alpha) +
  theme_ipsum(grid = F,axis=T,ticks=T,axis_text_size=15,axis_title_size = 15) +
  geom_hline(aes(yintercept = 0), colour="red")+ geom_smooth(method = "lm", se = FALSE, col="darkblue",linetype="dashed",lwd=0.4)+
  labs(x ="distance class index", y = "Mantel correlation")


DFTM_mantel<-tick.correlogM$mantel.res   
DFTM_mantel<-as.data.frame(DFTM_mantel)
class(DFTM_mantel)
tick.correlogM # determine the alpha values depending on the FDR corrected p values
DFTM_mantel$alpha<-c(1,0.1,0.1,0.1,0.1,0.1,0.85,0.1,0.1,0.85,0.1,0.1,0.1)
DFTM_mantel

  MantCorM<-DFTM_mantel %>%
  tail(100) %>%
  ggplot( aes(x=class.index, y=Mantel.cor)) +
  geom_line( color="grey") +
  geom_point(shape=21, color="black", fill="black", size=6, alpha=DFTM_mantel$alpha) +
  theme_ipsum(grid = F,axis=T,ticks=T,axis_text_size=15,axis_title_size = 15) +
  geom_hline(aes(yintercept = 0), colour="red")+ geom_smooth(method = "lm", se = FALSE, col="darkblue",linetype="dashed",lwd=0.4)+
  labs(x ="distance class index", y = "Mantel correlation")
  
  MantCorF + MantCorM  # joined plots to make it easier to insert in the manuscript
  
  
  # Pathogen diagnosis: GLM to assess the effect of "region" on infection
  
  Pathogen_ticks<-read.csv("~/Downloads/PathogenDx.csv", header = TRUE, sep = ",", row.names=1)
  Pathogen_ticks
  
  # replacing "inconclusive" diagnossis for "NA"
  
  Pathogen_ticks$E_chaff[Pathogen_ticks$E_chaff=="inconclusive"]<-NA
  Pathogen_ticks$E_chaff[Pathogen_ticks$E_chaff=="negative"]<-0
  Pathogen_ticks$E_chaff[Pathogen_ticks$E_chaff=="positive"]<-1
  
  Pathogen_ticks$E_ewingii[Pathogen_ticks$E_ewingii=="inconclusive"]<-NA
  Pathogen_ticks$E_ewingii[Pathogen_ticks$E_ewingii=="negative"]<-0
  Pathogen_ticks$E_ewingii[Pathogen_ticks$E_ewingii=="positive"]<-1
  
  Pathogen_ticks$R_rick[Pathogen_ticks$E_R_rick=="inconclusive"]<-NA
  Pathogen_ticks$R_rick[Pathogen_ticks$R_rick=="negative"]<-0
  Pathogen_ticks$R_rick[Pathogen_ticks$R_rick=="positive"]<-1
  
  Pathogen_ticks$A_phag[Pathogen_ticks$A_phag=="inconclusive"]<-NA
  Pathogen_ticks$A_phag[Pathogen_ticks$A_phag=="negative"]<-0
  Pathogen_ticks$A_phag[Pathogen_ticks$A_phag=="positive"]<-1
  Pathogen_ticks
  
  class(Pathogen_ticks)
  str(Pathogen_ticks)
  Pathogen_ticks$E_chaff<-as.numeric(Pathogen_ticks$E_chaff)
  Pathogen_ticks$E_ewingii<-as.numeric(Pathogen_ticks$E_ewingii)
  Pathogen_ticks$R_rick<-as.numeric(Pathogen_ticks$R_rick)
  Pathogen_ticks$A_phag<-as.numeric(Pathogen_ticks$A_phag)
  Pathogen_ticks$Region<-as.factor(Pathogen_ticks$Region)
  str(Pathogen_ticks)
  
  # Is E. chaffensis infection driven by region?
  # assuming normality
  GLM_tickPath_Echaff<-glm(E_chaff ~Region, data=Pathogen_ticks,family=binomial(link = "logit"))
  GLM_tickPath_Echaff
  plot(GLM_tickPath_Echaff)
  anova(GLM_tickPath_Echaff,test = "Chisq" )
  # p> 0.05 stil we will run the pothoc test for illustration
  summary(GLM_tickPath_Echaff)
  TukeyHSD(aov(GLM_tickPath_Echaff), "Region")
  
  # non-parametric test
  kruskal.test(E_chaff ~Region, data=Pathogen_ticks)
  # p>0.05 no need for Dunn test
  
  
  # Is E. ewingii infection driven by region?
  # assuming normality
  GLM_tickPath_Eewii<-glm(E_ewingii ~Region, data=Pathogen_ticks,family=binomial(link = "logit"))
  GLM_tickPath_Eewii
  plot(GLM_tickPath_Eewii)
  anova(GLM_tickPath_Eewii,test = "Chisq" )
  # p> 0.05 stil we will run the pothoc test for illustration
  summary(GLM_tickPath_Eewii)
  TukeyHSD(aov(GLM_tickPath_Eewii), "Region")
  
  # non-parametric test
  kruskal.test(E_ewingii ~Region, data=Pathogen_ticks)
  # p>0.05 no need for Dunn test
  
  
  # Is R. rickettsii infection driven by region?
  # assuming normality
  GLM_tickPath_Rri<-glm(R_rick ~Region, data=Pathogen_ticks,family=binomial(link = "logit"))
  GLM_tickPath_Rri
  plot(GLM_tickPath_Rri)
  anova(GLM_tickPath_Rri,test = "Chisq" )
  # p> 0.05 stil we will run the pothoc test for illustration
  summary(GLM_tickPath_Rri)
  TukeyHSD(aov(GLM_tickPath_Rri), "Region")
  
  # non-parametric test
  kruskal.test(R_rick ~Region, data=Pathogen_ticks)
  # p>0.05 no need for Dunn test
  
  
  # Is A. phagocytophilum infection driven by region?
  # assuming normality
  GLM_tickPath_Ap<-glm(A_phag ~Region, data=Pathogen_ticks,family=binomial(link = "logit"))
  GLM_tickPath_Ap
  plot(GLM_tickPath_Ap)
  anova(GLM_tickPath_Ap,test = "Chisq" )
  # p< 0.05 
  summary(GLM_tickPath_Ap)
  TukeyHSD(aov(GLM_tickPath_Ap), "Region")
  
  # non-parametric test
  kruskal.test(A_phag ~Region, data=Pathogen_ticks)
  # p<0.05 
  dunnTest(A_phag ~Region,
           data = Pathogen_ticks,
           method = "none")
  # no signficant differences between regions
  
  # Alpha diversity GLM analysis and box plot
  
  tick <- read.table("AlphaDivTicks.csv", sep=",", header=T)
  attach(tick)
  
  y <- glm(log(Observed) ~ Sex * Region, family = "gaussian",data=tick)
  plot(y)
  #fit generally good, some outliers
  
  y1 <- glm(log(Observed) ~ Sex + Region, family = "gaussian",data=tick)
  anova(y,y1, test="Chisq")
  #interaction not significant
  plot(y1)
  summary(y1)
  anova(y1, test="Chisq")
  # Analysis of Deviance Table
  # Model: gaussian, link: identity
  # Response: log(Observed)
  # Terms added sequentially (first to last)
  # 
  #         Df Deviance Resid. Df Resid. Dev  Pr(>Chi)    
  # NULL                     188    112.570              
  # Sex     1   13.862       187     98.708 1.351e-08 ***
  # Region  3   19.637       184     79.071 6.582e-10 ***
  #   ---
  # Signif. codes:  0 '***' 0.001 '**' 0.01 '*' 0.05 '.' 0.1 ' ' 1
  
  TukeyHSD(aov(y1), "Region")
  # Tukey multiple comparisons of means
  # 95% family-wise confidence level
  # Fit: aov(formula = y1)
  # 
  # $Region
  # diff        lwr       upr     p adj
  # MW-HC -0.3074068 -0.8203506 0.2055369 0.4077315
  # NE-HC  0.1799953 -0.1796971 0.5396877 0.5657488
  # W-HC   0.9788452  0.5789126 1.3787778 0.0000000
  # NE-MW  0.4874021 -0.1022631 1.0770673 0.1435051
  # W-MW   1.2862521  0.6712135 1.9012907 0.0000011
  # W-NE   0.7988500  0.3043403 1.2933596 0.0002530
  
  x <- glm(Simpson ~ Sex * Region, family = "quasibinomial",data=tick)
  plot(x)
  #fit generally good, some outliers
  
  x1 <- glm(Simpson ~ Sex + Region, family = "quasibinomial",data=tick)
  anova(x,x1, test="Chisq")
  #interaction not significant
  plot(x1)
  summary(x1)
  anova(x1, test="Chisq")
  # Analysis of Deviance Table
  # Model: quasibinomial, link: logit
  # Response: Simpson
  # Terms added sequentially (first to last)
  # 
  # Df Deviance Resid. Df Resid. Dev  Pr(>Chi)    
  # NULL                     188     52.569              
  # Sex     1  20.6001       187     31.969 < 2.2e-16 ***
  # Region  3   2.4027       184     29.566  0.002338 ** 
  summary(x1)
  TukeyHSD(aov(x1), "Region")
  # Tukey multiple comparisons of means
  # 95% family-wise confidence level
  # Fit: aov(formula = x1)
  # $Region
  # diff         lwr        upr     p adj
  # MW-HC -0.08938429 -0.22205869 0.04329011 0.3026856
  # NE-HC  0.10570734  0.01267186 0.19874283 0.0189076
  # W-HC   0.04012977 -0.06331396 0.14357350 0.7462250
  # NE-MW  0.19509163  0.04257301 0.34761024 0.0060011
  # W-MW   0.12951406 -0.02956747 0.28859559 0.1534960
  # W-NE  -0.06557757 -0.19348393 0.06232879 0.5455199
  
  
  par(mfrow=c(2,2))
  boxplot(log(Observed)~Region, col=c("white", "#9370DB" ,"#FFD700", "#FF0000"),ylab="Log(Observed ASVs)", cex.lab=1.2)
  boxplot(log(Observed)~Sex, ylab="Log(Observed ASVs)", cex.lab=1.2)
  
  
  boxplot(Simpson~Region, col=c("white", "#9370DB" ,"#FFD700", "#FF0000"), ylab="Gini-Simpson Index", cex.lab=1.2)
  boxplot(Simpson~Sex, ylab="Gini-Simpson Index", cex.lab=1.2)
  
  
  
  
  
